# A Lightweight Deep Learning Framework for Fast, Real-Time Super-Resolution Fluctuation Imaging

**DOI:** 10.1101/2025.06.05.658028

**Authors:** Miyase Tekpınar, Jelle Komen, Hana Valenta, Ran Huo, Klarinda de Zwaan, Peter Dedecker, Nergis Tomen, Kristin Grußmayer

**Affiliations:** Department of Bionanoscience and Kavli Institute of Nanoscience, TU Delft, Delft, Netherlands; Department of Intelligent Systems, TU Delft, Delft, Netherlands; Department of Chemistry, KU Leuven, Leuven, Belgium; Biomedical Engineering, Nanoscopy for Nanomedicine, TU Eindhoven, Eindhoven, Netherlands

**Keywords:** Real-time super-resolution imaging, low latency, AI, convolutional gated recurrent unit (convGRU), fluctuation microscopy, super-resolution optical fluctuation imaging, high-throughput imaging

## Abstract

Live-cell imaging captures dynamic cellular processes, yet many structures remain beyond the diffraction limit. Fluctuation-based super-resolution techniques overcome this limit by exploiting correlations in fluorescence blinking, but they typically require hundreds of frames and computationally intensive post-processing, prohibiting real-time imaging of fast cellular events. Recent deep learning approaches aim to increase the temporal resolution; however, many rely on extensive pre-processing or large, complex models that increase training cost and inference latency, preventing real-time deployment. To address this, we employ a light-weight recurrent neural network model, which integrates sequential low-resolution frames to extract spatio-temporally correlated signals. Our method is taylored for live-cell imaging under extreme signal-to-noise ratio conditions. It significantly improves temporal resolution by reducing the required number of frames down to as few as 8 frames while doubling the spatial resolution in an inference time below 30 ms. By combining simulation based training with an efficient network architecture, we introduce RESURF, a deep-learning based real-time super-resolution fluctuation imaging framework. We demonstrate that RESURF generalizes across different biological structures and can be readily adapted to various microscope setups using transfer learning. The accompanying dataset, comprising simulations and experiments across multiple subcellular structures and labeling strategies, establishes a benchmarking platform for fluctuation-based super-resolution techniques. RESURF offers a practical, low-latency deep-learning framework for high-throughput and real-time live-cell super-resolution imaging.

## Introduction

Live-cell fluorescence imaging captures dynamic cellular behaviors and aims to maximize both spatial and temporal resolution while minimizing sample damage [1], enabling advancements in fundamental cell biology. However, visualizing these cellular dynamics is constrained by the diffraction limit, which restricts the spatial resolution to around 200-300 nm. Super-resolution (SR) techniques can overcome this barrier and achieve higher spatial resolutions [2–4]. Single molecule blinking based super-resolution (SR) techniques use the stochastic switch between a bright “on-” state and dark “off-” state and can be performed with a wide-field microscope. Under sparse blinking conditions, single-molecule localization microscopy (SMLM) can be used to localize individual molecules and generate super-resolution images requiring many thousands of frames, usually corresponding to data acquisition over minutes to hours. As an alternative, fluctuation-based super-resolution techniques typically require only hundreds of frames to reconstruct a SR image as they can handle high densities of blinking fluorophores [5]. Widely used examples of fluctuation-based methods are super-resolution optical fluctuation imaging (SOFI) [5–7] and SRRF (Super-Resolution Radial Fluctuations) [8, 9].

SOFI leverages the correlation of blinking from a fluorophore over time and space and the independence of different fluorophores [6, 10]. Dertinger et al. showed that *n*^th^ order cumulants of a diffraction limited image sequence of blinking emitters result in a point spread function (PSF) raised to the power of *n*, which brings a 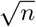-fold better resolution. This resolution enhancement can be further increased to n-fold in *n*^th^-order SOFI by applying linearization and deconvolution or Fourier reweighting after the cumulant calculation [11]. Another common technique is SRRF, which operates by computing gradient maps and applying a radiality transform that exploits the radial symmetry of the PSF. A final cumulant-based analysis of the image sequence produces a super-resolved, noise-suppressed image. The enhanced version, eSRRF, further assists users in selecting parameter settings that minimize the risk of artifacts [9]. Even-though the above-mentioned methods require shorter imaging time than SMLM, their limited speed restricts applications to slow cellular dynamics. In addition, SR image reconstruction is performed after image acquisition (off-line) and the computation typically takes seconds to minutes [12]. Therefore, these approaches are not suitable for real-time, simultaneous image acquisition and SR image reconstruction desirable for high-throughput and smart microscopy and in particular for live-cell imaging. These challenges have motivated the development of computational strategies aimed at enabling faster [10] and more efficient reconstruction [13, 14].

Deep learning has been extensively explored to increase image quality and imaging speed [15]. Most methods employ the U-Net architecture [16], a type of convolutional neural network (CNN), because of its high performance in image segmentation, denoising, and deconvolution[17–21]. U-Net is also used to accelerate SRRF imaging[21]. However, it restricts input image size and requires an interpolation step prior to U-Net inference, limiting its real-time deployment. In addition to CNNs, generative adversarial networks (GAN) can in principle be used for super-resolution image generation. SFSRM employs a GAN architecture called ESRGAN and a single-frame with an edge map as the network input [22]. In addition to edge map analysis, this method also requires a denoising step for low-SNR conditions prior to ESRGAN and the large size of the model results in a high computational latency of hundreds of ms in inference time, preventing real-time application.

Despite the growing number of studies that apply deep learning in microscopy, few AI-based methods focus on exploiting temporal correlations ([23, 24]). Recurrent neural networks (RNNs) are a notable deep-learning architecture to leverage temporal information, and are rarely used in microscopy. An example of an RNN-based SR technique, specifically a long-short-term memory (LSTM) network, is DBlink for video reconstruction from SMLM data [24]. By feeding localization maps into the LSTM, DBlink generates super-resolution videos at rates of up to 66 frames per second. However, this method remains constrained to SMLM sample conditions and relies on localization-based processing, restricting its use to off-line image analysis.

In this study, we offer a deep-learning framework to achieve real-time fluctuation-based super-resolution (RESURF) from few, ultra low-SNR microscopy images. RESURF enables real-time deployment by both reducing the imaging time, and by allowing a pre-processing free, fast inference time with only raw microscopy images as input. The reduced computational complexity and efficient training also result in reduced power consumption, crucial considering the growing use of AI in science and medicine.

We chose a deep-learning architecture based on a lightweight RNN architecture capable of leveraging both temporal and spatial information called multi-image super-resolution gated recurrent unit (MISRGRU [25]). As a target output we use second-order SOFI images to achieve a two-fold resolution improvement while limiting the computational demand of data preparation for training. Originally designed to predict single super-resolution images from multiple low-resolution images in remote sensing, MISRGRU utilizes a convolutional gated recurrent unit (convGRU), making it compatible with any fluctuation-based super-resolution technique. We show that it can be applied in bioimaging that in contrast operates under low-light conditions delivering high numerical aperture, diffraction-limited microscopy images. MISRGRU is flexible in that it supports training and inference with an arbitrary image size. By design, the output can be easily interpolated to any image size, allowing for straightforward training to produce n-fold larger output images, facilitating n-fold resolution improvement.

We used large synthetic data to create multiple foundation models, where we have full access to ground truth structures for evaluation. Our simulated dataset features microtubule-like structures with different SNR levels, dynamic out-of-focus background, and a realistic sCMOS camera noise model. We tested different input frame counts for the network to determine the model performance and the temporal resolution. Compared to a hundred or more frames typically required for second-order SOFI, our models achieved a two-fold spatial resolution improvement for very low-SNR conditions by using as few as 8 input frames. To allow real-time deployment, we optimized the size of the network and achieved an inference time less than 30 ms. We also showed the generalization of the model to different patterns using both experimental and simulated data. Furthermore, by sharing our simulations and experimental dataset, including multiple subcellular structures and labeling strategies, we offer our community a benchmarking platform for fluctuation-based microscopy methods.

Next, we demonstrate that we can optimize the foundation models via transfer learning with a small dataset for a range of live-cell and fixed-cell data labeled with different blinking fluorescent proteins and organic dyes. Using realistic simulations with the parameters close to experimental data for retraining, the network generalized to different structures and out-performed SOFI and eSRRF. In addition, we demonstrate that foundation models can learn different microscopy configurations from limited experimental data. This allows users to have flexibility in training strategies that can be conveniently adapted to facilitate dynamic and high-throughput super-resolution experiments.

## Results and discussion

### Enabling ultrafast and real-time super-resolution microscopy

SOFI typically requires several hundreds of frames (100-500) depending on fluorophore blinking characteristics and the signal-to-noise ratio (SNR) to reach its theoretical two-fold resolution enhancement and to produce high-SNR second-order SOFI reconstructions [26]. However, this frame count and the associated computational time limit SOFI’s ability to achieve high temporal resolution for real-time super-resolution sample screening. To address this limitation, we leverage deep learning to reduce the number of required input frames and computational time to enable real-time analysis, while maintaining image quality comparable to high-SNR SOFI reconstructions.

The deep-learning network architecture is based on the MISRGRU model, which comprises an encoder to extract feature representations of low-resolution frames, followed by a convolutional gated recurrent network (convGRU) based fusion layer to process the features sequentially to capture the correlations between input images. Global average pooling (GAP) combines them into a single representation, and a decoder reconstructs the high-resolution image by upsampling the fused feature maps. The input of the network consists of a time series of diffraction-limited frames, the target output being second-order SOFI reconstructions of the structures, as illustrated in Figure 1. To validate the network, we created a realistic synthetic dataset by generating structural patterns resembling microtubules randomly decorated with fluorophores at a certain density. We simulated fluorophore blinking dynamics, including on- and off-states, and incorporated an sCMOS camera noise model along with a non-stationary background to closely mimic experimental imaging conditions (see Methods). The dataset includes a range of SNR levels (photon number/frame: 40,60,80,100) and includes varying off-times (600, 1200, 2000 ms), while maintaining a constant average on-time of 10 ms to mimic gentle, low SNR live-cell conditions with fluorescence proteins labels. To our knowledge, this is the first large public dataset of fluctuation microscopy simulations intended to be used for training and benchmarking new methods. For generating SOFI target images, we used 100 diffraction-limited frames to compute second-order cross-cumulants, followed by linearization and Richardson-Lucy deconvolution for post-processing [11]. Following an approach similar to Chen et al., we trained separate models using 8, 15, 20, and 25 input frames to assess the network’s ability to reconstruct super-resolved images under different temporal constraints[21]. We also tested both mean-absolute error (L1) and Fourier loss (F) and showed that Fourier loss performs better in reaching higher RSP scores for slow-blinking low-SNR conditions (Supplementary Figure 1).

**Figure 1.**
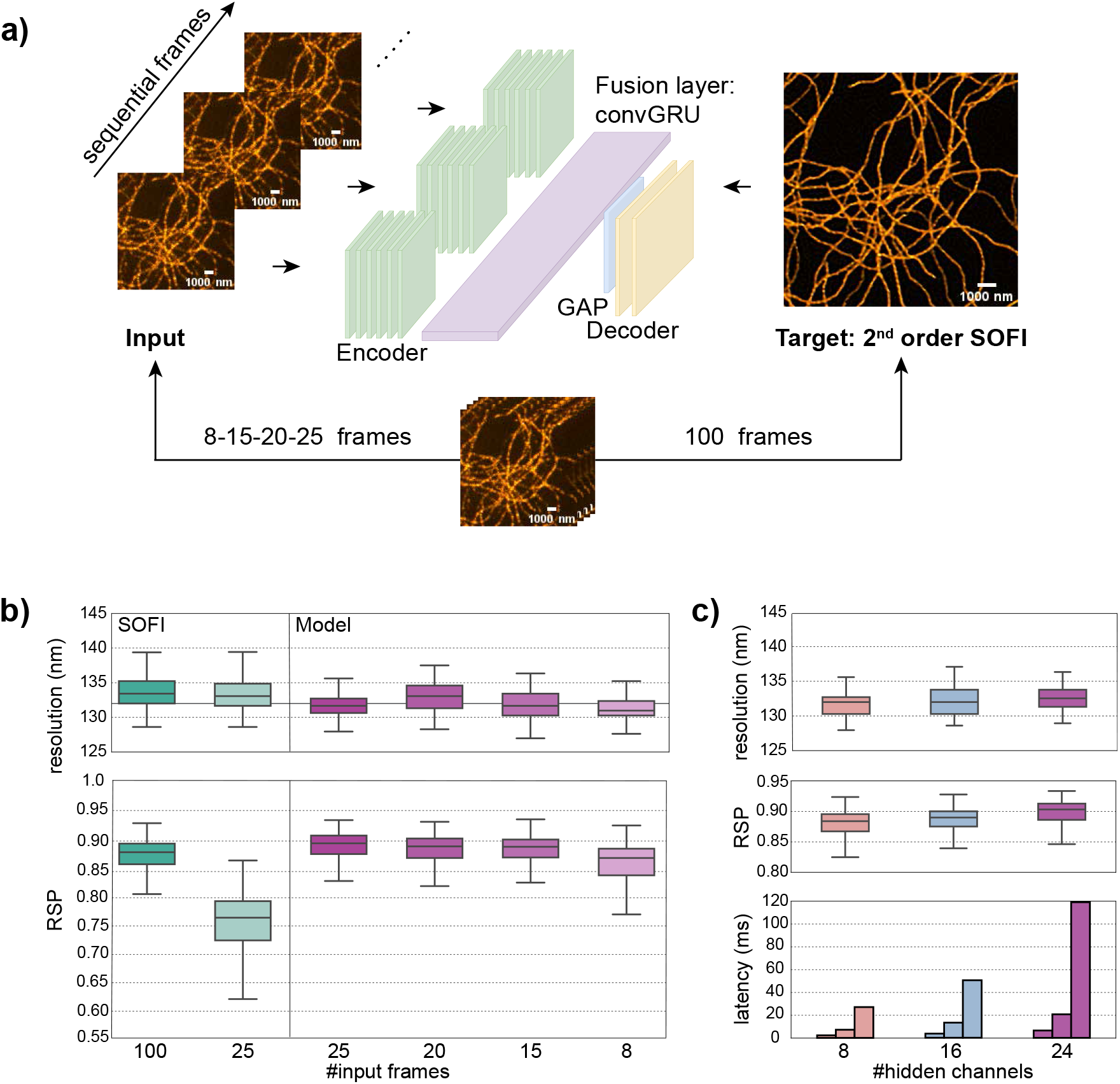
The network architecture and optimization for RESURF. **a.** The network architecture (MISRGRU) and training strategy. It is trained with simulated diffraction limited image sequences and corresponding second order SOFI reconstructions. Multiple models (foundation models) are trained by using different number of input frames (8,15,20,25) and, different number of hidden channels. **b**. Spatial resolution estimated via image decorrelation analysis and resolution-scaled Pearson correlation coefficient for SOFI (green) and the models (purple) using different numbers of input frames. Each boxplot depicts the median value (center line), 25th and 75th percentile (box edges), and the 1.5 IQR (interquartile range) (whiskers). The line at 132 nm marks the half-resolution of the widefield average image. **c**. Performance evaluation of models configured with different numbers of hidden channels: 8 (red), 16 (blue), and 24 (purple), all using 20 frames input. Latency is the average inference time over 500 repetitions (with TensorRT optimization): from left to right, each bar represents a different field of view (FOV) size: 128×128, 256×256, and 512×512 pixels. All models shown in this figure are trained by using F loss. The comparison of different loss functions can be found in Supplementary Figure 1. Scale bar: 1 µm.

We determined the performance of the models with different input frames by applying a set of established evaluation metrics, as described in more details in the Performance evaluation part of the Methods section. We estimated the resolution of predicted super-resolution (SR) images using decorrelation analysis, a widely adopted technique in super-resolution microscopy for resolution quantification [27]. We measured the similarity between the predicted and ground-truth (GT) or wide-field (WF) images using resolution-scaled Pearson correlation coefficient(RSP) and resolution-scaled error (RSE), a standard method in SR imaging for quantitative comparison of image quality [28, 29]

In figure 1b, we show the resolution and RSP scores of the models with different number of inputs and compare it to the SOFI method. All models approximate the theoretical two-fold spatial resolution improvement of second-order SOFI. The RSP scores reveal that all models surpass SOFI in preserving structural integrity. While the Pearson correlation coefficient slightly decreases with fewer input frames, we find that the 8-frame model offers a favorable trade-off between temporal resolution and spatial accuracy for highdensity samples. We optimized the network to limit computational latency and enable real-time deployment. We decreased the size of the model by systematically reducing the number of hidden channels while keeping the performance comparable to that of larger models. In figure 1c, we show that the RSP and resolution scores change only slightly for the reduced model size and the latency can be reduced down to 27 ms for an image size of 512×512 pixels and 20 input frames. The 8-frame model can offer higher temporal resolution and computational speed with only 11 ms inference time. Despite the inherent complexity of sequential data processing in recurrent neural networks, our optimized MISRGRU models achieved a significantly lower end-to-end latency of less than 30 ms. This represents an approximate 400-fold acceleration in super-resolution reconstruction compared to the SOFI method. These findings highlight the strong potential of MISRGRU for real-time, live-cell super-resolution imaging. We further compare our deep-learning framework with other approaches [21, 30], previously used for super-resolution acceleration, in Table 1. RESURF offers a lightweight, fast network model, and a computationally manageable simulation-based training pipeline to enable accessible deployment for biological research.

**Table 1:** Comparison among different deep-learning based super-resolution methods using different architectures. Input frames are the required low-resolution images for the input of the network. DL-eSRRF uses eSRRF images and SFSRM uses rendered localization microscopy image as a target. Pre-processing column shows the required steps before the inference of the model. Training samples includes each structure type with the number of image patches used during training. Parameters are the number of parameters to be trained for each model. Time shows inference latency of each model. It is benchmarked by using GPU, NVIDIA RTX A5000. For this table, we did not use any optimization such as TensorRT to speed up the inference.

| Method | Architecture | Input frames | Pre-processing | Training sample (size) | Parameters | Time (ms) |
| --- | --- | --- | --- | --- | --- | --- |
| DL-eSRRF [21] | U-NET | 5 | interpolation | microtubules (1005)<br>CCPs (727) | 31 M | 66 |
| SFSRM [30] | ESRGAN | 1 | edge map,<br>denoising | synthetic random emitters (1000)<br>microtubules (300)<br>vesicles (600) | 33 M | 732 |
| RESURF | MISRGRU-hc24<br>MISRGRU-hc8 | 8,20 | not required | synthetic microtubules (2000) | 131 K<br>15 K | 65, 158<br>23, 57 |

Visual examples of reconstructions under varying SNR levels and blinking conditions are provided in Figure 2. As expected, under the slowest blinking condition (Figure 2a), Model-8f misses some structures, simply because of the limited number of emitter switching to on-state in this short period of time. As we simulated the images with a Gaussian PSF and with FWHM of 220 nm, we first calculated the resolution of noise-free wide-field images (260 nm) and assigned the half of this value (130 nm) for the two-fold resolution improvement goal. However, image decorrelation analysis can be affected by the noise and background levels. Therefore, in Figure 2b, we see very high resolution values in wide-field images of slow blinking conditions with worse SNR and SBR compared to fast-blinking conditions. SOFI reconstructions using 100 frames fail to achieve a two-fold resolution improvement under low SNR and slow blinking conditions. On the other hand, Model-20f and Model-8f maintain the resolution improvement under different conditions. This showed that by being trained with a simulation dataset varied in SNR levels, the network learned to extract only the meaningful, correlated signal between the subsequent frames but not the uncorrelated noise. In Figure 2c, we tested the resolution-scaled Pearson correlation (RSP) scores of super-resolution reconstruction by taking both ground-truth and averaged wide-field frames as a reference in two separate analysis. The difference between the two analysis showed us that stationary background pixels in wide-field images degrade the RSP scores if the SR reconstruction performs a good background elimination. Wide-field images have very poor SNR levels under slow blinking low SNR conditions. As a result, the RSP estimations using wide-field images resulted in lower scores than the ones using ground-truth as a reference. In the latter case, we can see the high performance of both models under extreme noise conditions (Figure 2c) compared to SOFI, more aligned with the images shown in Figure2a. This offers a practical benefit for real-time super-resolution fluctuation (RESURF) live-cell imaging by tolerating lower laser intensities, thereby reducing phototoxicity and extending image acquisition durations.

**Figure 2.**
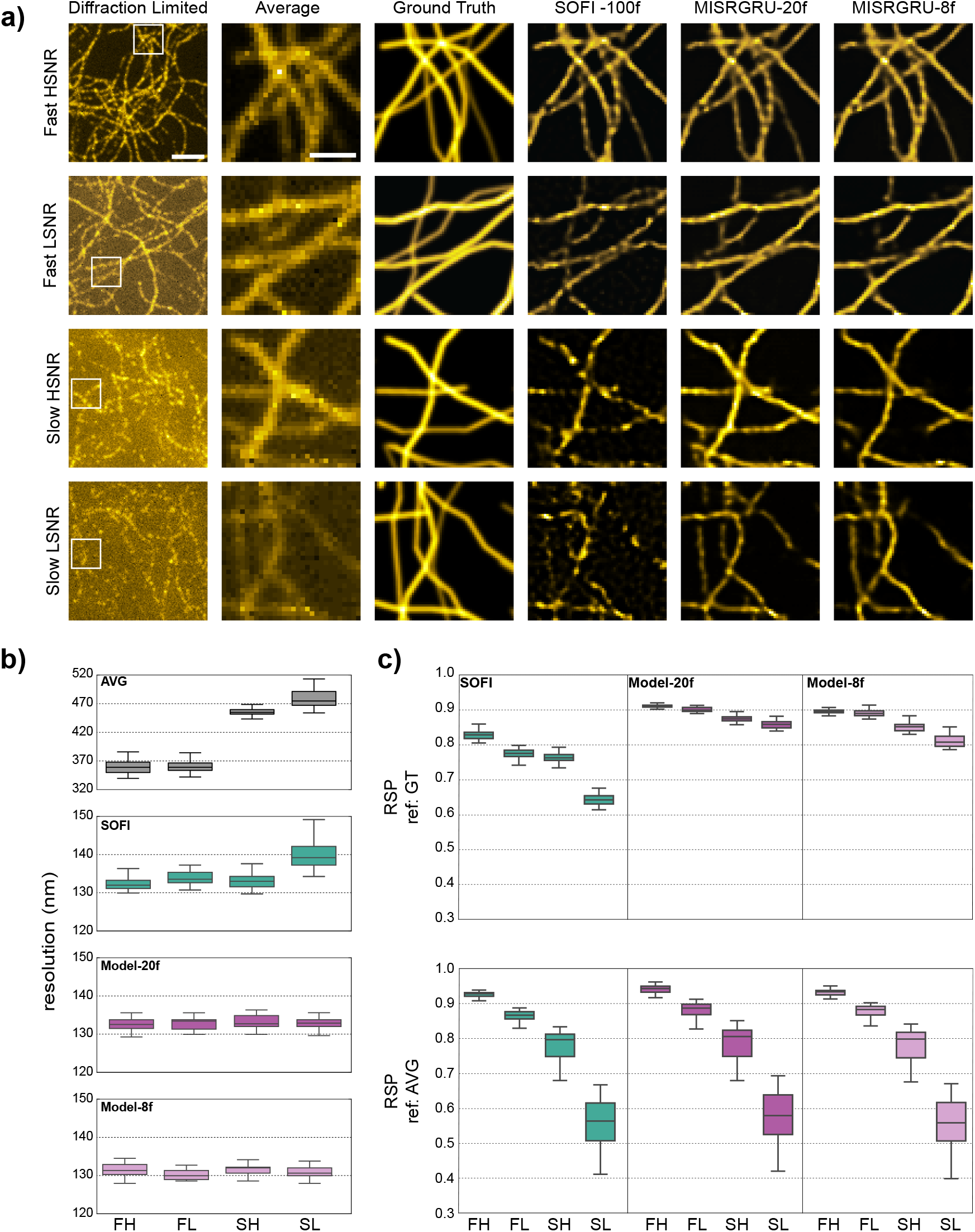
Performance of foundation models with simulations. **a.**Qualitative comparison of models and SOFI. Each row corresponds to a different SNR (photons/frame): 100(high-SNR) or 40 (low-SNR), and and blinking conditions (on time/off time in ms): 10/600 (fast) or 10/2400 (slow). From left to right: a single diffraction-limited input frame (128×128 pixels or 12.8×12.8 *µm*), the average of cropped area (30×30 px), the ground truth image demonstrating two-fold resolution enhancement, second-order SOFI reconstructions using 100 frames, and Model-20f and Model-8f. **b, c**. Corresponded resolution estimations and RSP scores of averaged wide-field, SOFI and the models for different conditions. RSP analysis is repeated for two different reference image: ground-truth (GT), and averaged wide-field (AVG) FH: Fast high SNR, FL: Fast low SNR, SH: Slow high SNR, SL: Slow low SNR. Pixel size in AVG: 100nm, Scale bars: 3 µm(diffraction limited), 1 µm (average).

In Supplementary Figure 1, we showed the performance of the network for the SNR conditions in and out of range of the training set. Models achieve generalization to different, higher and extremely low SNR levels which were not seen during training. Another model for higher SNR levels than the tested conditions can be easily retrained with a small dataset (see Transfer learning with small experimental dataset). In Supplementary Figure 5a, we evaluated the dependence of the model on the size of the training data set and showed that the model converges to the minimum training error already around the training set size of 2000. In addition, we also tested the model’s performance for different combinations of training datasets (k-fold cross validation analysis) and showed that models have high consistency, showing very similar scores in RSP and resolution analysis (Supplementary Figure 5c). Furthermore, we tested the performance of the models for different image sizes and demonstrated that model performance does not degrade for increasing image sizes, even though it is only trained using images with a size of 128×128 pixels (Supplementary Figure 6). This offers a widely-applicable method for a large variety of typical imaging conditions using modern scientific cameras without additional effort for retraining of the model.

### RESURF models generalize from simulations to different structures in live cells labelled by fluorescence proteins

Considering the dynamic features and poor SNR conditions of live-cell experiments, collecting a dataset for training and processing the data to create super-resolution target images can be very challenging. Therefore, we generated a new synthetic dataset closely mimicking our experiments with different photoswitchable fluorescence proteins (ffDronpa and SkylanS) as explained in. We used the specific, known parameters of the live-cell experiments to simulate their blinking kinetics and the full-width-half-maximum for the Gaussian-shaped point-spread-function obtained from our experimental data. A more detailed list of simulation parameters and network evaluation with simulations are provided in Supplementary Figure 2. As the SNR and SBR levels of the simulations were extremely low, 100 frames were not enough to create high SNR SOFI images as target for training. To mitigate the risk that the network could learn background pixels containing deconvolution artifacts caused by SOFI analysis, we applied additional filtering on SOFI images by using a wavelet-based background and noise subtraction (WBNS) [31].

In Figure 3 and 4, we show the performance of RESURF in fast-bleaching live-cell measurements by testing it with different intracellular structures and with different photoswitchable fluorescence proteins. As no ground-truth (GT) image is available in this setting, we employed resolution-scaled Pearson correlation (RSP) to assess structural fidelity and resolution-scaled error (RSE) to show the difference between the reference image and resolution scaled version of the super-resolution image [28]. As previously mentioned for Figure 2, RSP analysis is sensitive to background pixels. Considering that the majority of the field-of-view in most microscopy experiments will belong to background pixels, we repeated the RSP and RSE analysis by using averaged wide-field (AVG) and post-processed wide-field (AVG-WBNS) as reference images. We compared the model results with SOFI images generated by 100 frames, SOFI images after post-processing with WBNS (SOFI-WBNS) and eSRRF images generated by 20 frames. Because of the higher background levels in this new dataset, we additionally incorporated L1 loss together with Fourier loss (Model-FL) during training and compared it to the model using only Fourier loss (Model-F) (see section Network and training scheme).

**Figure 3.**
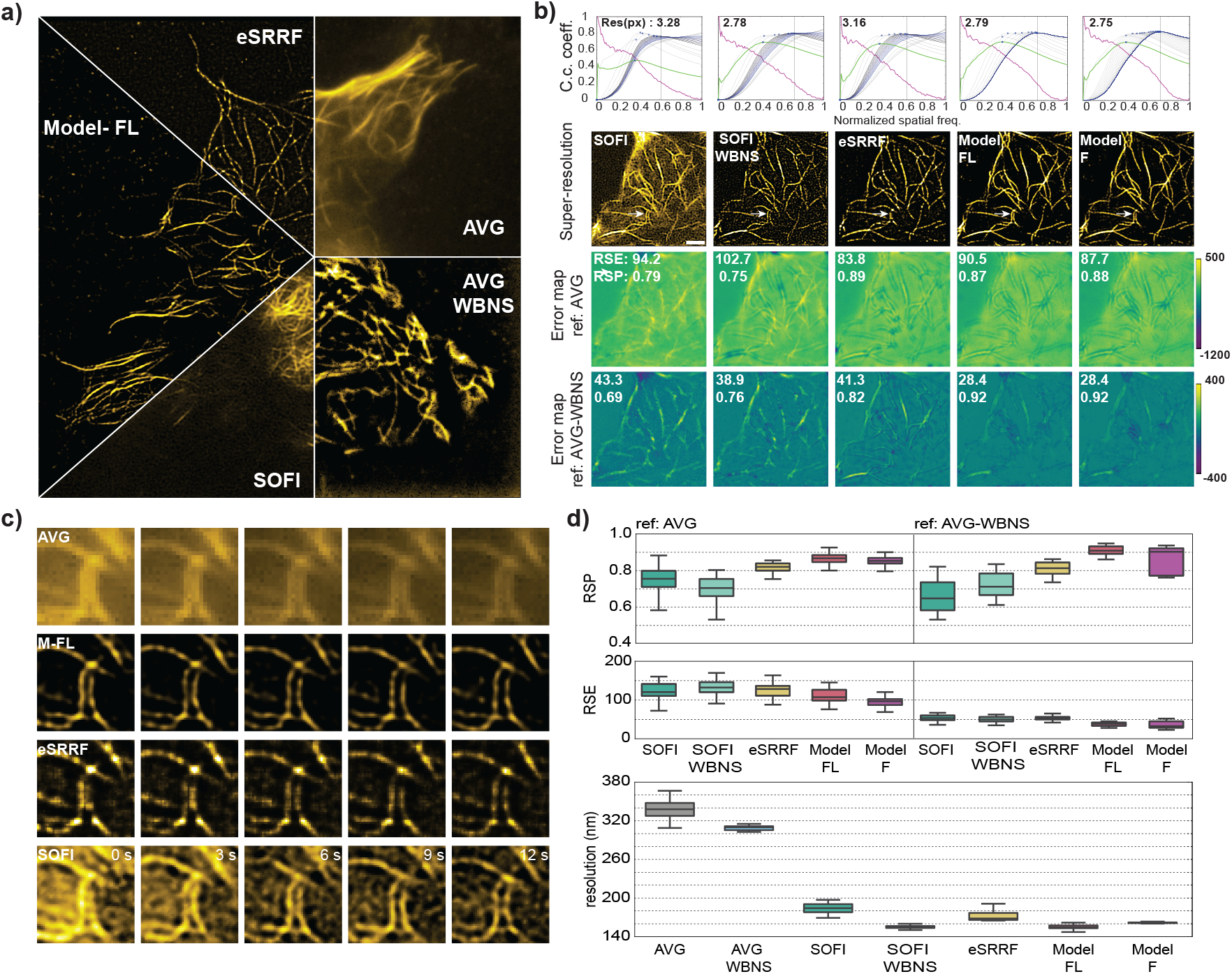
RESURF reconstructs SR images with high temporal resolution in a fast-bleaching live-sample. **a.**Qualitative comparison of SR reconstruction of microtubule structures using Model-FL, SOFI and eSRRF (HeLa cell, MAP4-ffDronpa). **b**. Influence of background on image decorrelation analysis and resolution-scaled error (RSE) for cropped regions of 128×128 pixels. First row shows decorrelation analysis for resolution estimation. Following SR reconstructions by different method RSE estimations for two reference images:(i) averaged projection of wide-field frames (AVG), (ii) background subtracted AVG images using wavelet-based noise and background subtraction (AVG-WBNS) **c**. SR reconstruction from a small dynamic region of 30×30 pixels (1.5×1.5 *µm*). **d**. Evaluation scores for resolution estimation, RSP and RSE by using AVG and AVG-WBNS as reference images. Each box plot is created by using 50 random crops of 128×128 from the large FOV in (a). Model-F: trained with Fourier loss, Model-FL: trained with combination of Fourier and L1 loss. Pixel size in AVG: 108 nm, Scale bars: 1.6 µm (b), 2 µm (c)

**Figure 4.**
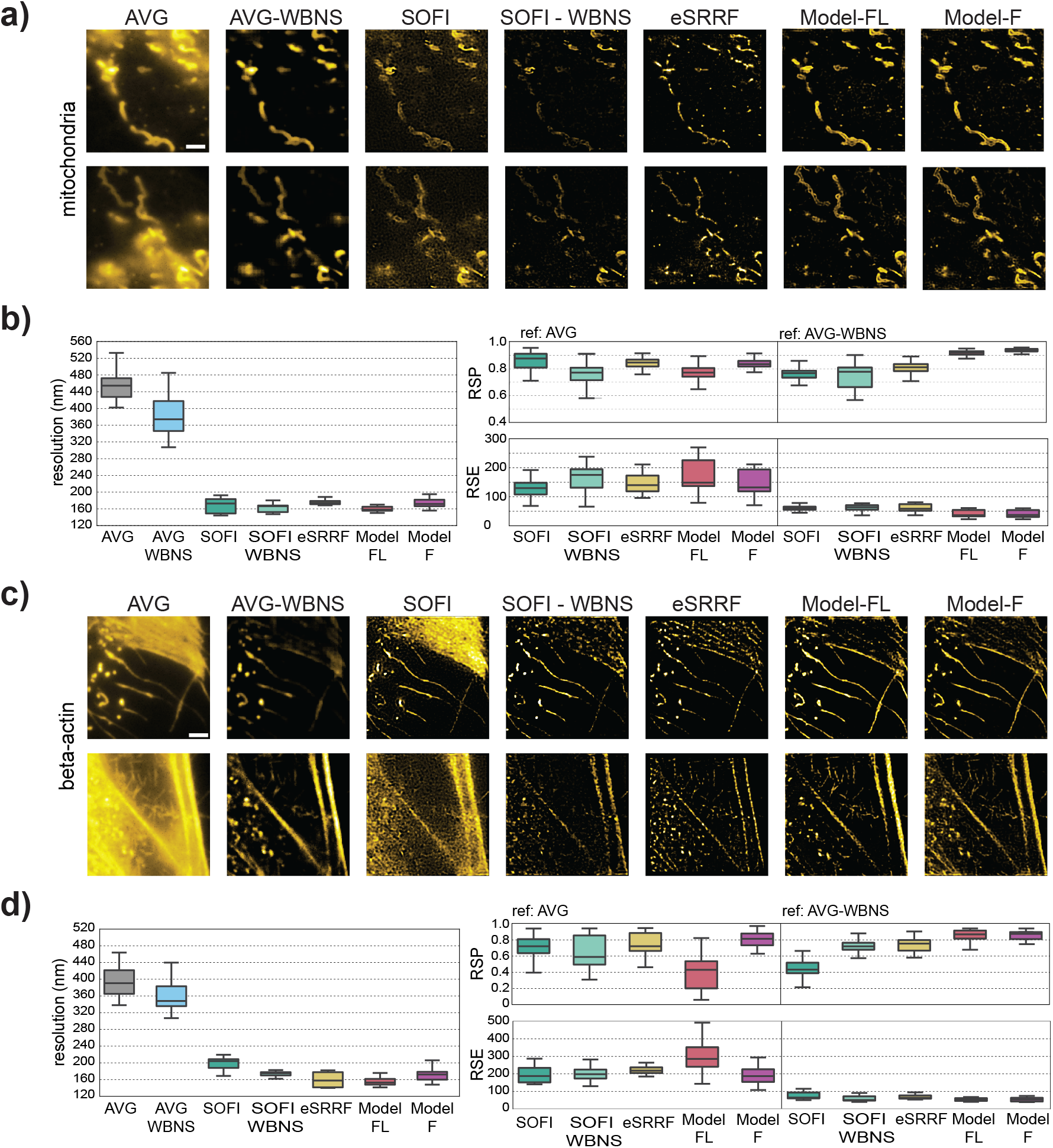
Generalization of RESURF to different intracellular structures. **a,b** Qualitative comparison of our models, SOFI and eSRRF for a region of 128×128 pixels for mitochondria (DAKAP-SkylanS) (a) and actin (SkylanS-betaActin) in HeLa cells (c). **(b,d.)** Correspondong evaluation scores for resolution estimation, RSP and RSE by using AVG and AVG-WBNS as reference images. Model-F: trained with Fourier loss, Model-FL: trained with combination of Fourier and L1 loss.Pixel size: 108 nm. Scale bars: 2 µm

In Figure 3b, we evaluate the super-resolution (SR) reconstructions by different methods. In the top row, we show the image decorrelation analysis graph for each SR image. The high background level in SOFI reconstruction influences image decorrelation analysis by resulting in a worse resolution estimation than SOFI-WBNS. eSRRF can eliminate background better than SOFI, however it cannot reach the resolution level that SOFI-WBNS and the models achieve. Two very close microtubules that clearly demonstrate this difference are indicated by white arrows and also shown in Figure 3c. Following the SR images, resolution-scaled error (RSE) maps for two different reference images are given in the last two rows, together with the average RSE score and resolution-scaled Pearson correlation coefficient (RSP). The RSP and pixel errors from RSE analysis are higher in the background when the reference image has a higher background and the super-resolution reconstruction eliminates the background. This effect can be clearly seen by comparing the RSE maps of SOFI and SOFI-WBNS for different reference images. Therefore, using AVG-WBNS as a reference gives more representative results for our models. The complete movie of the cell in Figure 3a is shown in Supplementary video 1.

As can be seen in Figure 4a and c, our models trained only on microtubule structures generalize well to mitochondria and actin structures. This is confirmed by evaluation scores almost identical compared to microtubule structures. We also note that Model-F and Model-FL perform similarly where high frequency spatial components such as microtubule and mitochondria images are needed to be separated from low-frequency spatial components such as large cellular membrane or background signal. As Model-FL is trained with both Fourier and pixel-level loss function L1 while targeting background-subtracted SOFI images, it does not recognize the low-frequency spatial components(4c,d) and thus performs better in background elimination. Overall, both models achieved generalization to different structures and did not show any sign of memorizing the patterns.

For experiments with high-background conditions, eSRRF serves as a complementary benchmarking method. SOFI and eSRRF have distinct advantages and limitations. SOFI typically requires larger number of frames and additional post-processing to perform well under high background levels, whereas eSRRF is known to achieve more effective background suppression with fewer frames. However, eSRRF can generate artifacts by collapsing closely spaced structures into one another [32]. This effect is particularly evident in the mitochondria images shown in Figure 4.

To achieve better generalization and avoid hallucinations, we followed several strategies. First, in addition to different SNR levels, we simulated different background levels and random emitters in different axial positions. Second, we optimized the SOFI targets to represent the information in the diffraction limited input images. And lastly, we kept the emitter density levels moderate and the number of input frames low, thus there are more random single emitters on the focal plane to avoid the structural bias. In addition to successful models, we also showed the counter examples in Supplementary Figure 3. When we trained the model to target the SOFI results using high SNR and an order of magnitude more frames, the model showed more hallucinations in the background. Next, we trained the network by increasing the emitter density. The resulting model learned to connect random emitters to create filamentous structures, showing high structural bias.

In summary, we calibrated our training dataset to have a large variety of SNR and background levels, while avoiding complete structures in SOFI targets and eliminating background via post-processing when necessary. As a result of our training strategy and the sequential processing ability in the convGRU layer, the models learn to extract the correlation between low-resolution images rather than recognizing similar patterns to the ones in the training dataset.

### Applying RESURF models on public datasets with different fluorescent labels

We further investigated the generalization of models trained with synthetic data to different microscopes by using public datasets. As we did not vary the PSF width in our simulations, our models require retraining to perform well in very different optical configurations (see section Transfer learning with small experimental dataset). Therefore we only selected datasets with a FWHM close to our synthetic images (between 2.5 and 3.2 pixels). First, we tested a dataset which uses self-blinking dyes to label microtubule structures with our model trained for live-cell conditions [33]. This public dataset was aimed at higher orders of SOFI analysis, using statistics from many frames with slower and more sparse blinking conditions than the datasets we tested in previous sections. As there are not enough emitters activated during 20-frames to reconstruct a complete structure, we averaged consecutive model predictions. By using either 100 or 8000 frames in total (by averaging 5 and 400 SR predictions), we compared them with the corresponding SOFI images. The conditions of the experiment were ideal for SOFI and both model and SOFI reconstructions showed comparable results (Fig. 5). Next, we used two datasets originally acquired for the deep-learning based super-resolution technique called SFSRM[30]. SFSRM used single frames to reach SMLM level resolutions. While the method can reach high temporal resolution, as stated in the study, this is an ill-posed problem and comes with strict limitations. Especially in low SNR levels, the method requires denoising prior to inference, making its application more complicated for live-cell conditions. The data set includes highly dynamic two intracellular structures labeled by fluorescent proteins, requiring high temporal information. Therefore we used 8-frames model and we did not apply any pre-processing before the model inference. We reconstructed a super-resolution movie of microtubule-vesicle interactions during vesicle transport and mitochondrial fission at endoplasmic reticulum contact site (Fig. 6). We showed that RESURF model can be used for wide-range of SNR, background and blinking conditions and is applicable to datasets from different microscopes with similar optics without retraining.

**Figure 5.**
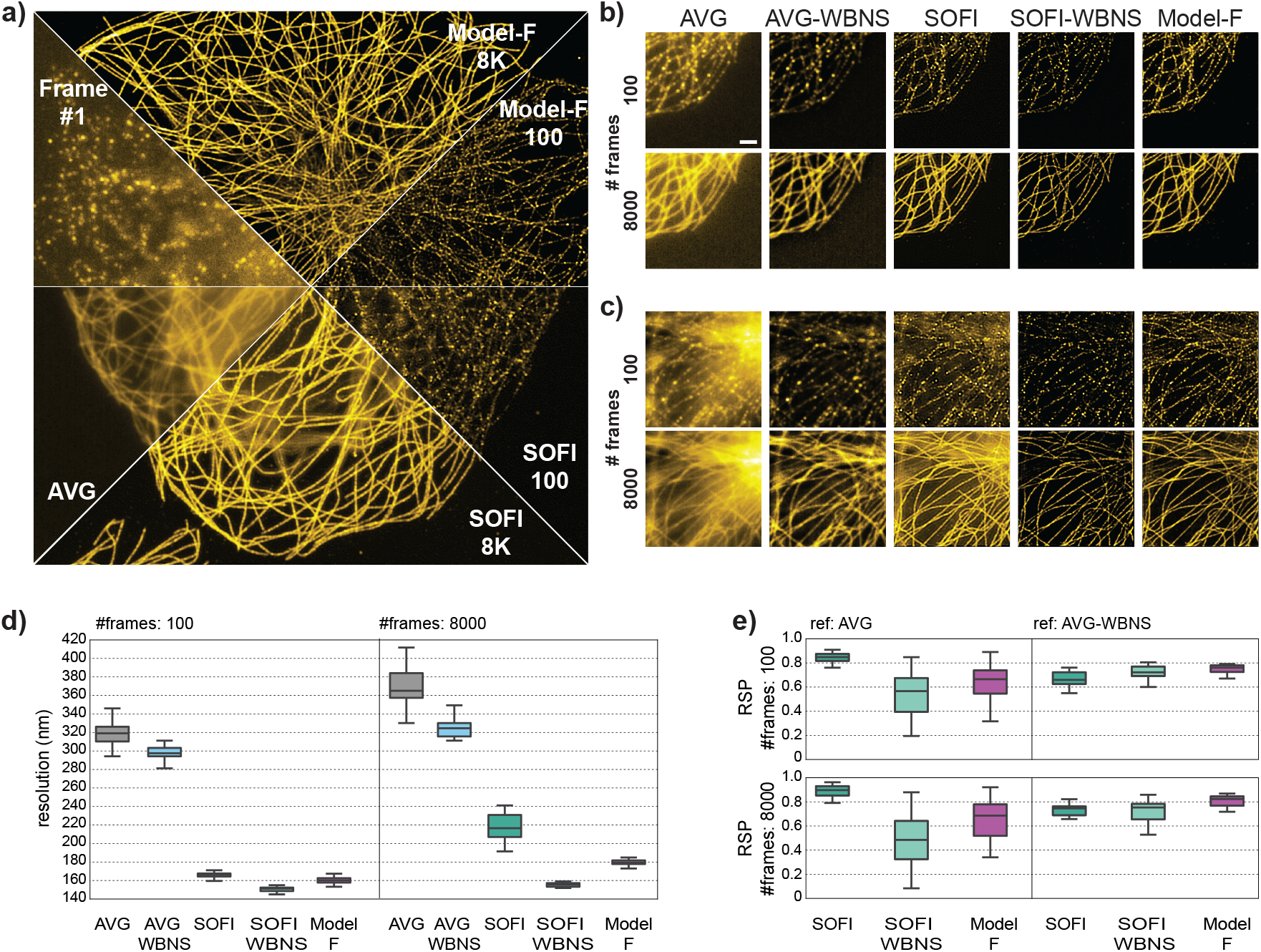
Generalization of RESURF to slow-blinking samples tested by a public SOFI dataset expressing microtubules and labelled by self-blinking dyes[33]. **a,b,c.** COS-7 cell, immunostained for microtubules with Abberior FLIP 565. Qualitative comparison of SR reconstructions using 100 or 8000 frames by our models, SOFI and eSRRF. Small regions of 128×128 pixels are cropped from a high structural density (c) and a low structural density (b) area. **d,e**. Quantitative comparisons via resolution estimation and resolution-scaled Pearson correlation (RSP). SOFI-8000 is the average of SOFI reconstructions using 500 frames. Model-F for 100 and 8000 frames are calculated by averaging the model reconstructions used by 20-frames inputs. Scale bars: 2 µm

**Figure 6.**
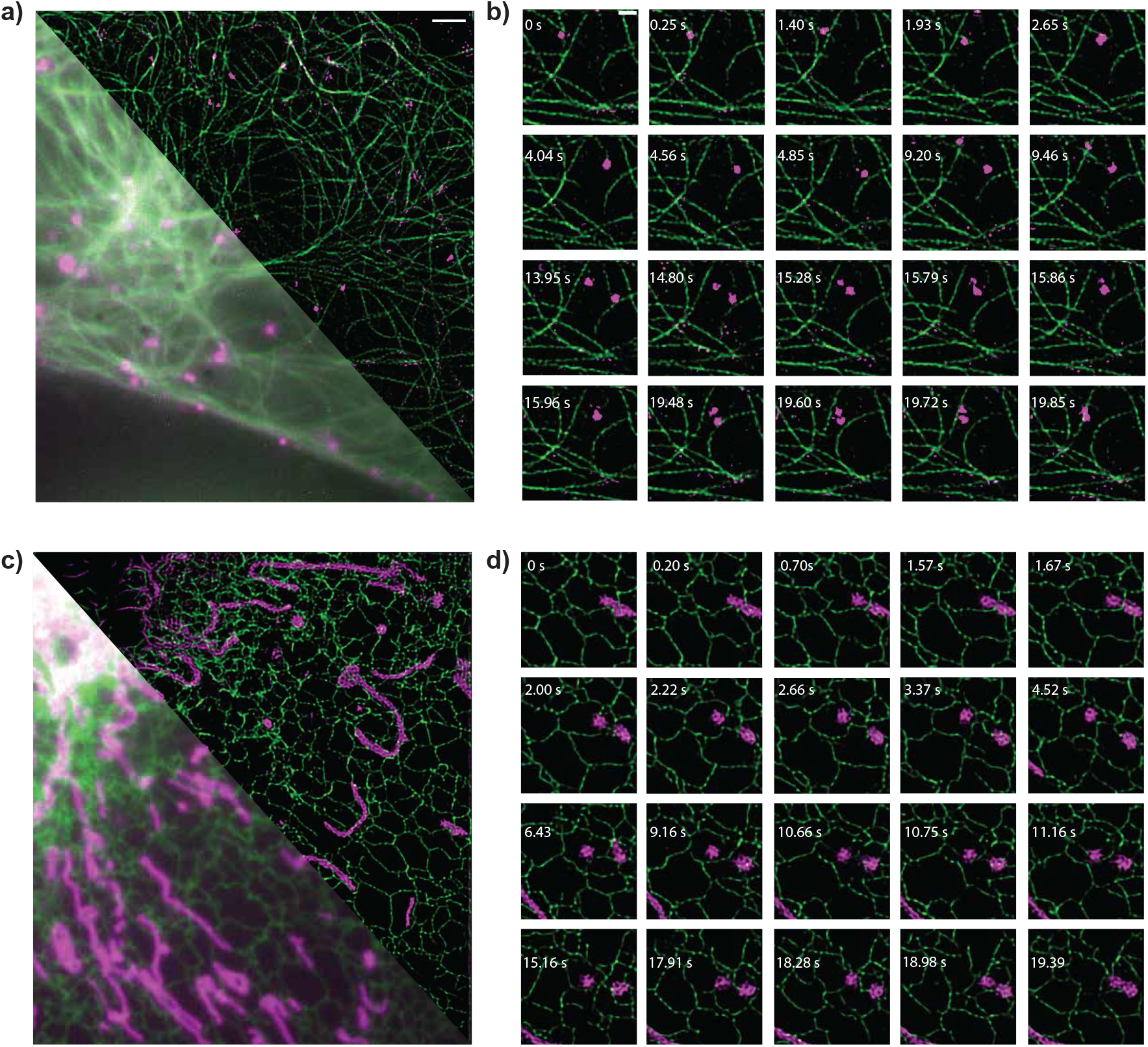
RESURF observation of dynamic biological processes in public dataset[30]. **a.** Two-color image of microtubules (green) and vesicles (magenta) in Beas2B cells. **b**. Example of vesicle transportation dynamics. **c**. Two-color image of endoplasmic reticulum (ER-green) and mitochondria (magenta) in Beas2B cells. **d**. Example of mitochondrial fission at ER-mito contact site. Wide-field images are averaged projection of first 8-frames and RESURF results are reconstructed by rolling a 8-frames window and using Model-F. Time-labels belong to the first frame in each window. Scale bars: 5 *µm* (a), 1 *µm* (b)

### Transfer learning with small experimental dataset

The previous models are trained only with simulations and focused on low and ultra low SNR conditions with a narrow PSF FWHM (around 2.7 pixels). We re-trained the network to achieve reliable SR prediction for high SNR DNA-PAINT experiments with a significantly larger PSF (FWHM of 3.8 pixels). We used transfer learning for the new model and trained it with experimental microtubule data acquired from fixed-cell samples. Therefore, we were able to evaluate the model’s performance for high-SNR applications and examine its consistency compared to models trained with simulations only. DNA-PAINT labeling can be used to tailor fluorescent blinking to achieve both SMLM and SOFI imaging [34]. As a test case, we chose a fast-blinking condition and as a result of abundant exchangeable bright emitters inside the DNA-PAINT buffer, it allows long-term high-SNR data acquisition. We showed that RESURF speeds up both data-acquisition and computation time, making it suitable for real-time high-throughput imaging.

Using pre-trained weights from synthetic data allows to leverage learned features, speeding up the training process and improving generalization, which is especially useful when constrained by limited microscopy data [24, 35, 36]. We trained the model using a dataset of only 500 patches of 128×128 pixels. We created ground-truth SOFI images by using 500 frames and trained the models using 20 frames input. As shown in Supplementary Figure 4 and for live-cell experiments, limiting the number of frames to be used for target SOFI reconstruction is a crucial step to prevent hallucinations and encourage generalization. Intuitively, using larger number of frames for SOFI could lead to better target images for training. However, it also brings a risk of hallucinations if the input images do not have the same emitters activated during 20 frames. This can encourage the model memorize the patterns instead of extracting temporal correlations. Therefore, although our dataset contains 10,000 frames of acquisition, we limit the target SOFI reconstructions to 500 frames to reduce the risk of hallucinations. The results are presented in Figure 7 and the model predictions are compared to both SOFI with 500 and 10,000 frames. Similar to the simulations, the models effectively approximates the filaments by doubling the resolution and shows high RSP score. This proves that RESURF is suitable for retraining with small datasets to learn different microscopy settings allowing easy adaptation and fast training for different experimental conditions.

**Figure 7.**
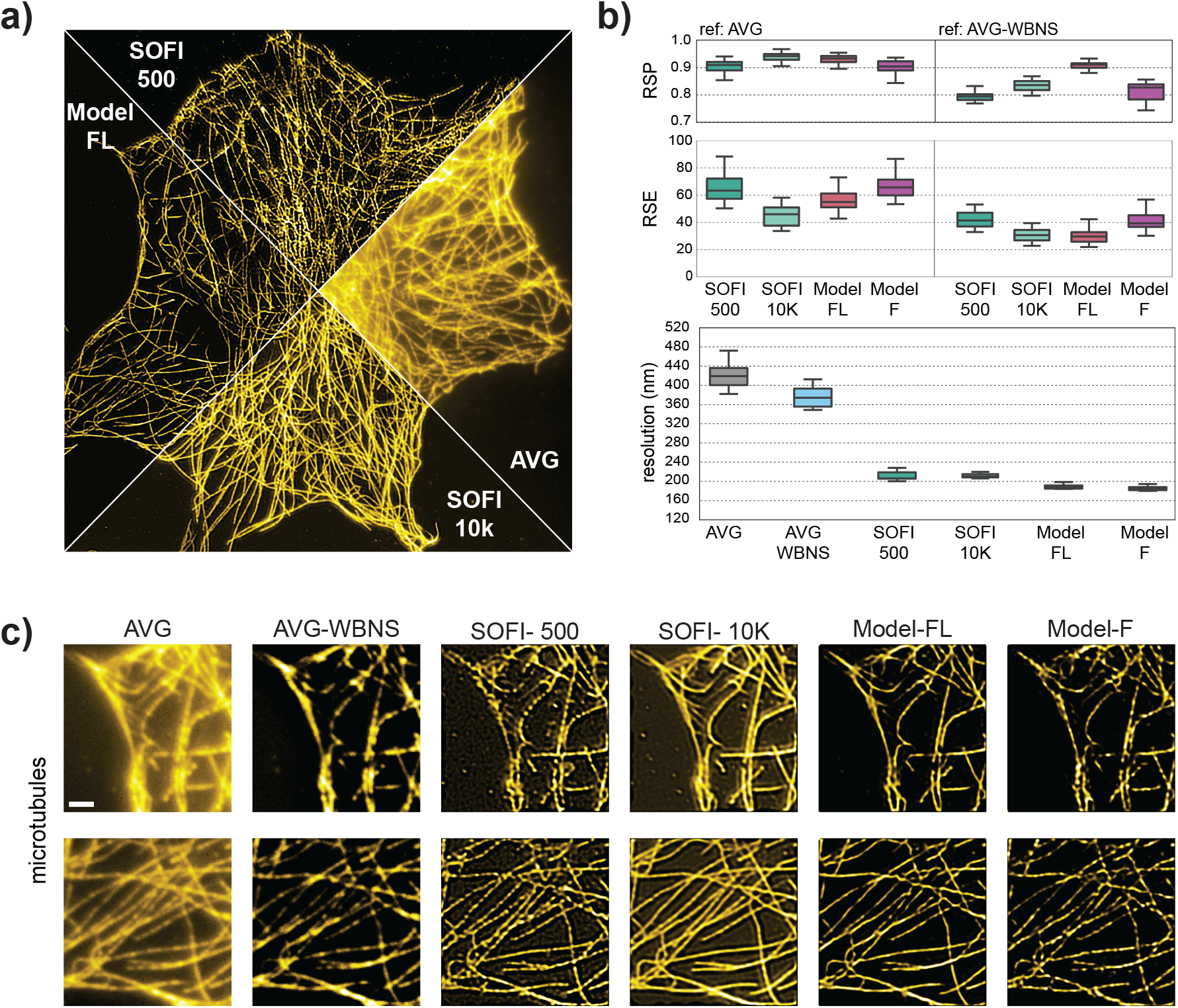
RESURF fine-tuned by small experimental dataset enables high-throughput SR imaging for fast-blinking DNA-PAINT samples. **a.**Super-resolution reconstructions and averaged wide-field image of microtubules in a COS7 cell, labeled by DNA-PAINT **b**. Corresponded evaluation scores of resolution estimation and resolution-scaled Pearson correlation (RSP) for two different reference images (average (AVG)and post-processed average (AVG-WBNS) of diffraction limited frames). **c**. Qualitative comparison of SR reconstructions by our models, SOFI with 500 and 10,000 frames for a cropped region of 128×128 pixels in wide-field. Averaged wide-field and model predictions are reconstructed by using 20 frames. Scale bar: 2 µm

## Discussion

We offer an effective, light-weight deep-learning solution for ultrafast super-resolution fluctuation imaging (RESURF). To utilize temporal information and reach a high computational speed, we adopted a deep-learning architecture with recurrent neural network called multi-image super-resolution gated recurrent unit (MISRGRU). We designed RESURF to enhance both spatial and temporal resolution while remaining adaptable to a range of imaging conditions. By the inherent ability of recurrent networks to process sequential data, our method could serve as a real-time alternative to any fluctuation-based super resolution imaging method. We also provide our simulation tool and dataset as a benchmarking platform for fluctuation based super-resolution techniques which can be used for both training and testing. We optimized our method with synthetic and experimental datasets to reconstruct second-order SOFI images with approximately 400-fold acceleration in processing time compared to conventional SOFI and using only 8 to 20 frames. RESURF allows forecasting super-resolution images of raw data during the ongoing experiment and reduces background artifacts while doubling the spatial resolution. We note that the latency of models benchmarked with a single GPU (NVIDIA RTX A5000) after TensorRT optimization could be further improved with more investment in hardware and parallelization. Although reducing the number of input frames enhances temporal resolution and increases the computational speed, this alone does not guarantee real-time deployment. Additional demands, such as pre-processing steps and model complexity, making other methods unsuitable [21, 30], are equally critical for a computationally manageable training pipeline and achieving real-time performance. Our focus was on developing an efficient, lightweight deep-learning pipeline that enables practical and accessible deployment. This has become increasingly important as artificial intelligence is progressively integrated in biomedical applications. Our models offer substantial reductions in computational cost, an important consideration to make deep learning accessible and reduce the carbon footprint.

Our model leverages temporal information to effectively capture correlated fluorophore blinking events, making it suitable for denser fluorophore labeling compared to SMLM. We observed that our approach achieves a favorable balance between spatial resolution, structural fidelity, and temporal performance. In principle, higher resolution improvement is also possible by using higher-order SOFI images as a target. Under extreme low signal-to-noise ratio (SNR) conditions, RESURF outperforms SOFI and eSRRF, suggesting its potential for live-cell imaging with reduced laser intensity, thereby minimizing photodamage and allowing for longer imaging sessions. In highly dynamic imaging scenarios, using a smaller number of input frames, in this case 8, reduces motion blur and enables the reconstruction of high-quality, high-frame-rate movies. Its real-time reconstruction performance allows on-the-fly segmentation, classification, or rare event detection in super-resolved images.

We first pre-trained the model only on synthetic datasets to create foundation models. This offers a great opportunity for live-cell experiments where collecting training data can be very challenging because of the poor SNR conditions and dynamic behavior of the sample. Next, we fine-tuned foundation models using a small number of data either from simulations or experiments. Leveraging pre-trained weights from synthetic data enabled the model to retain useful feature representations, thereby accelerating convergence and enhancing generalization. This is particularly valuable when only limited experimental microscopy data are available. Moreover, the models were able to generalize to different intracellular structures, including mitochondria and actin, as well as vesicles and endoplasmic reticulum acquired from a different microscopy system, as tested using public datasets. By optimizing the simulation conditions, SOFI target images resemble an incompletely sampled structure and do not have the full underlying ground truth [20]. Thus, our models can extract temporal correlation between input frames without structural bias.

For our foundational model, we focused on low and ultra-low SNR conditions. Furthermore, we preserved the original 16-bit dynamic range of the images and did not apply min–max normalization. This design choice was intended to maintain consistent brightness relationships between independent inputs during inference. However, this increases the model’s sensitivity to high-intensity inputs, which may result in saturated predictions. Mitigation is straightforward through transfer learning using a small dataset, here demonstrated for a high-SNR DNA-PAINT sample.

We optimized our training to eliminate background artifacts by post-processing target SOFI images (synthetic dataset) or using longer image sequences to reconstruct the SOFI image (experimental dataset). To further improve the robustness of the model, especially under challenging imaging conditions such as those encountered in live-cell cytosolic environments, several strategies can be pursued. One promising direction is enhancing the quality of target images used during training through advanced post-processing techniques, such as improved linearization and through sophisticated pre-processing methods. These improvements are critical only for ultra low-SNR regimes, where SOFI reconstructions tend to degrade in quality. Heterogeneous experimental conditions constitute another challenge for deep learning approaches. Multiple models could be trained for different background and SNR conditions instead of one global model and applied for different regions of the images. This approach has recently been successfully implemented for SMLM [37]. An alternative could be increasing the number of learnable parameters by increasing the number of channels and using complementary targets, instead of one SOFI image, to balance between missing structures and background artifacts. This could allow the model to learn more complex tasks. However, optimizing a larger model comes with the cost of higher computational latency and training complexity.

In summary, we present a lightweight recurrent neural network-based architecture tailored for real-time fluctuation super-resolution imaging (RESURF) within the microscopy domain. The proposed model is capable of reconstructing high-quality super-resolution images from as few as eight input frames, while minimizing background artifacts and achieving a two-fold spatial resolution enhancement near the theoretical limit. By leveraging pre-trained weights from synthetic datasets and fine-tuning with either synthetic or fixed-cell microscopy data, the network is well-suited for live-cell imaging scenarios. Thanks to its low latency and practical training workflow, the model holds strong potential for real-time sample screening and for enhancing spatio-temporal resolution during live experiments. Furthermore, fast super-resolution streaming enables high-throughput imaging and opportunities for smart microscopy.

## Methods

### Simulations

The MATLAB based SOFI simulation tool was adapted to more accurately represent noise and background effects, following the methodology described in [38]. To account for electronic noise, a gain and standard deviation map of the sCMOS camera used in our experimental setup was generated and incorporated into the simulation [39, 40]. In addition, a non-stationary background model was implemented in which background signals arise from out-of-focus blinking emitters. Inspired by the approach used in SACD [10], the distribution of emitters in the focal plane was geometrically shifted and transformed to generate a corresponding background distribution, which was subsequently convolved with a broader point spread function (PSF) to simulate out-of-focus blur. For the simulations prepared for live-samples, we added random emitters in focal plane and in some random axial planes outside focal plane.

The simulation begins by generating a binary mask of the target structure, which consists only filamentous patterns for training datasets of all the models presented in this study. We also created hollow mitochondria structures to test the the generalization of the model to different structures. These patterns are produced by modifying the pattern generation algorithms originally developed for the DBlink method and to study mitochondrial dysfunction [24, 41]. Emitters were randomly distributed across the simulation area at a density of 5000 fluorophores*/*µm^2^, and only those overlapping with the binary mask were retained. The spatial grid for emitter placement was set to 1280×1280 pixels with a resolution of 10 nm per pixel. Filament structures with thicknesses ranging from 10 nm to 40 nm were simulated.

The temporal blinking behavior of individual emitters was simulated using a continuous-time Markov process [38], resulting in high-resolution video sequences. The emitters have average on-time of 10 ms, and averaged off-times of 600 ms, 1200 ms, or 2400 ms to model different blinking speeds. Signal-to-noise ratio (SNR) was further controlled by adjusting the Ion value, which represents the average number of photons emitted per frame by each fluorophore. Ion values of 100, 80, 60, and 40 were used to reflect different SNR conditions, mimicking the photophysics of fluorescent proteins under low illumination.

Both in-focus and out-of-focus emitter planes were convolved with Gaussian PSFs, having full-width at half-maximum (FWHM) values of 220 nm and 2000 nm, respectively. The maximum background contribution per pixel was calculated based on the mean brightness of fluorophores in the focal plane, using a fixed ratio between in-focus and out-of-focus signals. The resulting noise-free image stack was then downsampled by a factor of 10. Finally, Poisson-distributed photon noise and sCMOS-specific electronic noise were added to each frame to produce the final simulated dataset.

These parameters are changed for fluorescence protein simulations to represent higher background and low SNR conditions as shown in Supplementary Figure 1 and 2. The on-time and off-time are adjusted for the fluorescence proteins and specifically for ffDronpa [42]. As a result, the on-time ratio happens to be faster than the simulations for the foundation models which could decrease the frequency of the appearance of random single emitters. Therefore we lowered the emitter density to 1000 fluorophores*/*µm^2^. As we used the foundation model to initialize the training, we only simulated 200 different microtubules patterns. Then, we varied the SNR and background levels by generating 10 different images for each pattern. Unlike previous simulations, we also added random emitters in the background, both in the focal plane and in different axial positions out of focus. In each 10-mers of simulation, we only added one high SNR condition and we maintained low SNR and high background conditions for the rest.

### Network and training scheme

Multi-image super-resolution (MISR) methods have been extensively studied in satellite imaging [43]. MISRGRU was first introduced in 2020 to the computer vision community to improve the quality of satellite imagery [25]. This model leverages multiple low-resolution images and an encoder-decoder framework to generate a super-resolved output. As illustrated in Figure 1, the architecture is composed of an encoder, a convolutional gated recurrent unit (convGRU) serving as a fusion layer, and a decoder. We inherited this architecture and modified to reduce size thus improve the latency by excluding the image registration layer and reducing the number of input channels from 64 to either 24 or 8. We also optimized the loss function by testing mean-absolute error as a spatial a spatial loss (*L*_1_), Fourier loss (*L*_ℱ_)[44]. During the training for the live-samples labeled by fluorescence proteins, we combined these two loss function to optimize structural integrity and resolution of predictions. We updated the weights for spatial and Fourier loss in very 20 epochs by using gradient normalization [45].

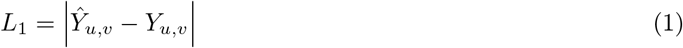

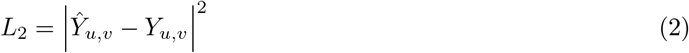

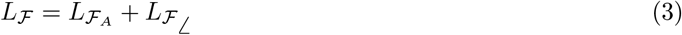

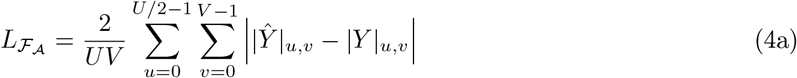

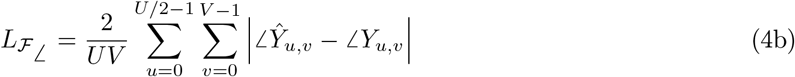

Here, both SR images *Ŷ* and the target image *Y* are transformed into the Fourier space by applying fast Fourier transform (FFT), where the absolute amplitude difference 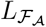 and absolute phase difference 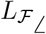 are calculated. Due to symmetry in the Fourier space (Hertimitian symmetry), only half of the spectral components is considered.

We trained the foundation model with simulated datasets of 128×128 pixel images. The dataset was split into training (2,000 samples), evaluation, and test sets (480 samples each). Model training was performed using the Adam optimizer with a learning rate (*η*) of 0.001 and momentum parameters *β*_1_ = 0.9 and *β*_2_ = 0.999. He Normal initialization was used for the model weights [46]. To mitigate overfitting, early stopping was applied with a maximum of 400 training epochs and a patience threshold of 10. A batch size of 10 was selected to balance GPU memory usage and computational efficiency. All training and testing were conducted on an NVIDIA GeForce RTX 3090 GPU with 24 GB of memory.

To extend the model to experimental data, we used the pre-trained weights from the foundation model as initialization of the models we trained for live-cell and DNA-PAINT measurements. To further reduce inference latency, we applied TensorRT acceleration to all models trained within the PyTorch framework [47]. This optimization included techniques such as layer fusion and precision calibration, including the use of lower-precision formats like fixed-point floating point, to maximize computational efficiency.

### Performance evaluation

We evaluate the performance of SOFI reconstructions and AI model generated predictions using three complementary methods. First, we estimate image resolution through image decorrelation analysis, a technique that operates on a single image and relies on partial phase correlation to determine spatial resolution [27]. Second, we employ the resolution-scaled Pearson correlation (RSP) metric to quantify the similarity between the super-resolution (SR) image and a reference image [48]. Prior to correlation computation, the SR image is adjusted to match the resolution of the reference image using an estimated resolution-scaling function (RSF), providing a more robust comparison than traditional metrics like structural similarity index (SSIM). For experimental data, the reference is the average projection of the image sequence (wide-field). As we noticed that the RSP score can under-represent the structural similarity in images with high-background, we also used a wavelet-based background subtraction method (WBNS) to create a background free alternative reference image [31]. We used 3 levels in wavelet decomposition to estimate the background and skipped the denoising part of the algorithm. In addition, for simulated test set, we also calculated the RSP scores between ground-truth and super-resolution predictions. Lastly, to visualize the pixel-level difference between reference image and the super-resolution prediction, we used resolution-scaled error map (RSE). Similar to RSP, absolute difference between pixel values are calculated after matching the resolution between two images and averaged to calculate RSE score. We noticed that the boundary pixels in model predictions for actin data (in Figure 4d) can sometimes show hot pixels. This is likely due to the extremely high signal levels in actin experiments, which is not represented in our training set. Therefore, we cropped the five boundary pixels in actin images and used the smaller, cropped region during the model evaluation with RSP and resolution analysis.

### Data analysis

#### SOFI

We reconstructed SOFI images using sofipackage based on cross-cumulant-based algorithm which is available from https://github.com/kgrussmayer/sofipackage. Following n-th order cross-cumulant analysis, images reach to resolution improvement of 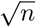-fold. Next, we apply linearization and Richardon-Lucy deconvolution as a post-processing [11, 26, 49]. To compensate the non-linearity in brightness, we either use linear-scaling or adaptive linearization. While the linear scaling takes the n-th root of the brightness in n-th order SOFI, adaptive linearization incorporates blinking properties to enable the effective use of the dynamic range. For Richardson-Lucy deconvolution (RLD), we used a gaussian kernel with the same FWHM we used for simulations and 10 iterations to reach two-fold resolution improvement. For foundation models, we use adaptive linearization and 2.2 pixels of FWHM for RLD. For high background simulations, imitating live-cell condition, we use linear-scaling and 2.7 pixels of FWHM for RLD and follow with a wavelet-based background and noise subtraction (WBNS) method to eliminate the background and ringing artifacts caused by deconvolution [31]. We used 3 levels decomposition for background estimation and 2 levels of decomposition for noise estimation.

For synthetic training datasets created for live-cell experiments, we followed WBNS method with a binary mask to eliminate the background pixels [50]. To create the binary mask, we applied Li-thresholding on the high-SNR versions in each 10-mers dataset.

#### eSRRF

We used the Fiji plugin of eSRRF to get guidance in finding the best parameters for radius (R) and sensitivity level (S), and operated the rest of the analysis using the Python tool, nanopyx [8, 29]. eSRRF has different types of temporal analysis. In all figures and evaluations, we used averaged eSRRF reconstructions with two factor interpolation.

### Sample preparation and data acquisition

#### DNA-PAINT imaging

Fast DNA-PAINT data used in Figure 7 were acquired using the method previously published [34]. Briefly, COS7 cells were cultured in DMEM with standard supplements, seeded on coverslips, and fixed. Microtubules in cells were stained firstly with primary antibodies against alpha-tubulin, followed by nanobodies conjugated with 5×R1 DAN-PAINT docking strands (sequence 5’-3’: TCCTCCTCCTCCTCCTCCT). Cells were imaged in buffer consisting of 20 nM R1 imager strands (sequence 5’-3’: AGGAGGA – Atto655), 5% (v/v) ethylene carbonate and 500 mM NaCl in PBS. Custom-built TIRF microscope was used for imaging at 638 nm excitation (2.5*kW/cm*^2^); 10,000 frames were recorded at 10 ms exposure using MicroManager. Full description of the nanobody production, labeling protocol and microscope specifications are available in the method section of [34].

#### Live-cell imaging

For live-cell imaging, HeLa cells were grown at 37^°^*C*, in 5% CO2 atmosphere in DMEM (ThermoFisher) complemented with 10% fetal bovine serum (FBS) and GlutaMAX (ThermoFisher). For imaging experiments, cells were seeded in 35-mm glass-bottom dishes (MatTek) and after 24 h, they were transiently transfected using FuGENE HD Transfection Reagent (Promega) according to the manufacturer’s protocol using following plasmids; MAP4-ffDronpa (created in the Dedecker Lab), DAKAP-SkylanS (created in the Dedecker Lab) and SkylanS-betaActin (purchased from Addgene). 24 hours after the transfection, the cells were washed with PBS (ThermoFisher) and then imaged in PBS on a Nikon Eclipse Ti-2 Inverted Microscope equipped with a 1.4 NA oil immersion objective (100 CFI Apochromat Total Internal Reflection Fluorescence) and a ZT405/488/561/640rpcv2 dichroic mirror with a ZET405/488/561/640 nm emission filter (both Chroma Technology). The imaging was performed always in TIRF mode using 488 nm and 405 nm lasers (Oxxius L6Cc laser combiner). Each measurement consisted of the acquisition of stacks of 500 or 1000 images. MAP4-ffDronpa and DAKAP-SkylanS are recorded at 30 ms and SkylanS-betaActin is recorded at 20 ms exposure.

## Acknowledgements

We thank the TU Delft Department of Bionanoscience for support and the TU Delft AI lab initiative for funding BIOlab. K.S.G was supported by an ERC grant (QScope, 101165129) funded by the European Union. Views and opinions expressed are however those of the author(s) only and do not necessarily reflect those of the European Union or the European Research Council Executive Agency. Neither the European Union nor the granting authority can be held responsible for them.

## Contributions

M.T, J.K and K.S.G conceived the idea. M.T designed the simulations. M.T and J.K designed the evaluation protocol with input from K.S.G and N.T. J.K tested MISGRU network and optimized them to create foundation models. M.T performed further analysis to evaluate the performance and improved the simulations and the network training for live-cell experiments and generalization of the models. M.T and J.K trained the model for transfer learning with experimental data. M.T analyzed public datasets to test generalization. H.V designed and performed the live-cell experiments. B.K.Z and R.H. designed and performed the DNA-paint experiments. M.T wrote the manuscript. K.S.G, N.T and P.D reviewed the manuscript. N.T supervised the deep network implementation and evaluation. K.S.G supervised the microscopy experiments, data analysis and network evaluation.

